# A combination of RBD and NTD neutralizing antibodies limits the generation of SARS-CoV-2 spike neutralization-escape mutants

**DOI:** 10.1101/2021.06.10.447999

**Authors:** Denise Haslwanter, M. Eugenia Dieterle, Anna Z. Wec, Cecilia M. O’Brien, Mrunal Sakharkar, Catalina Florez, Karen Tong, C. Garrett Rappazzo, Gorka Lasso, Olivia Vergnolle, Ariel S. Wirchnianski, Robert H. Bortz, Ethan Laudermilch, J. Maximilian Fels, Amanda Mengotto, Ryan J. Malonis, George I. Georgiev, Jose A. Quiroz, Daniel Wrapp, Nianshuang Wang, Kathryn E. Dye, Jason Barnhill, John M. Dye, Jason S. McLellan, Johanna P. Daily, Jonathan R. Lai, Andrew S. Herbert, Laura M. Walker, Kartik Chandran, Rohit K. Jangra

**Author notes:** Corresponding authors (R.K.J.), (K.C.). These authors made equivalent contributions (Denise Haslwanter, M. Eugenia Dieterle).

## Abstract

Most known SARS-CoV-2 neutralizing antibodies (nAbs), including those approved by the FDA for emergency use, inhibit viral infection by targeting the receptor-binding domain (RBD) of the spike (S) protein. Variants of concern (VOC) carrying mutations in the RBD or other regions of S reduce the effectiveness of many nAbs and vaccines by evading neutralization. Therefore, therapies that are less susceptible to resistance are urgently needed. Here, we characterized the memory B-cell repertoire of COVID-19 convalescent donors and analyzed their RBD and non-RBD nAbs. We found that many of the non-RBD-targeting nAbs were specific to the N-terminal domain (NTD). Using neutralization assays with authentic SARS-CoV-2 and a recombinant vesicular stomatitis virus carrying SARS-CoV-2 S protein (rVSV-SARS2), we defined a panel of potent RBD and NTD nAbs. Next, we used a combination of neutralization-escape rVSV-SARS2 mutants and a yeast display library of RBD mutants to map their epitopes. The most potent RBD nAb competed with hACE2 binding and targeted an epitope that includes residue F490. The most potent NTD nAb epitope included Y145, K150 and W152. As seen with some of the natural VOC, the neutralization potencies of COVID-19 convalescent sera were reduced by 4-16-fold against rVSV-SARS2 bearing Y145D, K150E or W152R spike mutations. Moreover, we found that combining RBD and NTD nAbs modestly enhanced their neutralization potential. Notably, the same combination of RBD and NTD nAbs limited the development of neutralization-escape mutants *in vitro*, suggesting such a strategy may have higher efficacy and utility for mitigating the emergence of VOC.

**Importance:** The US FDA has issued emergency use authorizations (EUAs) for multiple investigational monoclonal antibody (mAb) therapies for the treatment of mild to moderate COVID-19. These mAb therapeutics are solely targeting the receptor binding domain of the SARS-CoV-2 spike protein. However, the N-terminal domain of the spike protein also carries crucial neutralizing epitopes. Here, we show that key mutations in the N-terminal domain can reduce the neutralizing capacity of convalescent COVID-19 sera. We report that a combination of two neutralizing antibodies targeting the receptor binding and N-terminal domains may have higher efficacy and is beneficial to combat the emergence of virus variants.

## Introduction

Severe acute respiratory syndrome coronavirus-2 (SARS-CoV-2), is a member of the family *Coronaviridae*, and the causative agent of the ongoing coronavirus disease 2019 (COVID-19) pandemic (1). Over 171 million cases have been officially diagnosed since its first emergence and >3.6 million people have succumbed to disease (2). Public health measures, along with rapid vaccine development have helped slow the pandemic in some countries. Moreover, small molecule inhibitors, antibody-based therapeutics, and convalescent plasma from COVID-19 convalescents have received emergency use authorizations (EUAs) (3). Recently, multiple virus variants of concern (VOC), some carrying neutralizing antibody (nAb)-resistant mutations that are associated with increased transmission and fatality rates, have emerged (4). The availability of multiple therapeutic approaches especially for people who cannot get vaccinated is essential. There is thus an urgent need to develop therapeutics, especially ones that limit the emergence of neutralization-resistant variants or are more efficient against them as they can help save lives while vaccines are being deployed.

SARS-CoV-2 entry into host cells is mediated by the transmembrane spike (S) glycoprotein, which forms trimeric spikes protruding from the viral surface (5). Each monomer, 180-200 kDa in size, comprises S1 and S2 subunits that are generated by post-translational cleavage by the host enzyme furin. The S1 subunit is composed of two domains, an N-terminal domain (NTD) and a C-terminal domain (CTD). The CTD functions as the receptor-binding domain (RBD) for the entry receptor, human angiotensin-converting enzyme 2 (hACE2) (6, 7). The role of the NTD for SARS-CoV-2 is unclear, but it has been proposed in other coronaviruses to play roles in recognizing specific sugar moieties during attachment and regulating the prefusion-to-postfusion transition of the S protein (8–10). The S2 subunit is composed of the fusion peptide, heptad repeats 1 and 2, a transmembrane domain and a cytoplasmic tail. Aided by hACE2-binding and host cathepsin- and/or transmembrane protease serine 2 (TMPRSS2)-mediated proteolytic processing, S2 undergoes extensive conformational rearrangement to insert its fusion peptide into the host membrane and mediate the fusion of host and viral membranes (6, 7).

The S protein is the major target of nAbs, the production of which is a key correlate of protection following virus infection and vaccination (11–14). Due to their potential to interfere with hACE2 interaction and to efficiently neutralize virus infection, RBD-specific antibodies have been the main focus of human monoclonal antibody (mAb)-based therapeutics (13, 15–20). We recently described the memory B-cell repertoire of a convalescent SARS donor and isolated multiple RBD-specific antibodies that neutralize and protect against SARS-CoV, SARS-CoV-2 and WIV1 viruses (19, 20). Since that time, multiple RBD-targeting mAbs have received emergency use authorizations by the US FDA. However, the widespread circulation of nAb-resistant variants has led to the withdrawal of EUAs for some nAb monotherapies (21) highlighting the need to develop combination-nAb therapies that can treat SARS-CoV-2 variants and reduce the probability of mutational escape. In fact, a few of the VOC carry mutations in some of the major neutralizing epitopes in the RBD as well as the NTD (22).

Recently, multiple NTD mAbs with potent neutralizing activity have been described (17, 23–28). As combinations of mAbs targeting distinct epitopes and mechanisms of action have been successfully used against other viruses (29, 30), cocktails of RBD and NTD mAbs have been proposed and recently showed promise against SARS-CoV-2 *in vitro* and *in vivo* (23, 24).

To evaluate the effect of combining nAbs targeting the RBD and the NTD, here, we mined the memory B-cell repertoires of four convalescent COVID-19 donors with high serum neutralization and spike-specific antibody titers. By sorting spike-reactive single B-cells, we isolated 163 mAbs targeting S. Further, we evaluated their neutralization capacity against authentic SARS-CoV-2 and a self-replicating vesicular stomatitis virus carrying SARS-CoV-2 S protein (rVSV-SARS2) (31). We downselected the top RBD-and NTD-targeting neutralizers and used multiple approaches to map their epitopes. As described recently (32, 33), we observed that neutralization-escape rVSV-SARS2 mutants of the NTD-targeting mAb were resistant to neutralization by COVID-19 convalescent donor sera, suggesting that natural variants in the NTD could, at least in part, escape the antibody response. Here, we show that a combination of the NTD-and RBD-targeting mAbs neutralized virus more efficiently and limited the emergence of neutralization-escape spike mutants, underscoring the utility of combination therapy.

## Results

### SARS-CoV-2 induces robust and diverse memory B-cell response in convalescent patients

To characterize the B-cell responses induced by SARS-CoV-2 infection, we sampled peripheral blood mononuclear cells (PBMCs) from four adult patients (EMC 5, 9, 10 and 15) at the Montefiore Medical Center in the Bronx (**E**instein-**M**ontefiore **C**OVID-19). They were all previously healthy individuals who developed mild COVID-19. SARS-CoV-2 infection in their nasopharynx was confirmed by a positive RT-qPCR test in the first week of March 2020. All four patients had convalescent blood drawn to collect serum and PBMCs at least two weeks after all symptoms had resolved on March 31, 2020 (**Fig. S1A**). Serum samples of all four donors displayed reciprocal serum neutralization IC^50^ titers of >118 against rVSV-SARS2.

For each donor, we single-cell sorted SARS-CoV-2 S-reactive class-switched (CD19^+^IgM^−^IgD^−^) B-cells, which ranged in frequency between 0.6 and 1.2% across the donors (**Fig. S1B-C**). Index-sorting analysis revealed that the S-specific B-cells were predominantly IgG^+^, and the majority expressed the classical memory B-cell marker CD27 (41-76%) (**Fig. 1A**). Additionally, 44-67% expressed the activation/proliferation marker CD71 (**Fig. 1B**), consistent with the early time post-infection. Antibodies from all four donors showed similar levels of somatic hypermutation, as evidenced by the median number of nucleotide substitutions in the heavy-chain variable region that are consistent with the early time point post infection (range 1-3) (**Fig. 1C**). Although VH germline genes such as VH3-30 and VH3-30-3 were over-represented in all individuals as has been seen previously (27), less than 5% of clones were derived from clonally expanded lineages (**Fig. 1D-E**). Altogether, these results indicate a robust and diverse early memory B-cell response to SARS-CoV-2 infection in each donor and are consistent with previous studies (34–36).

**FIG 1.**
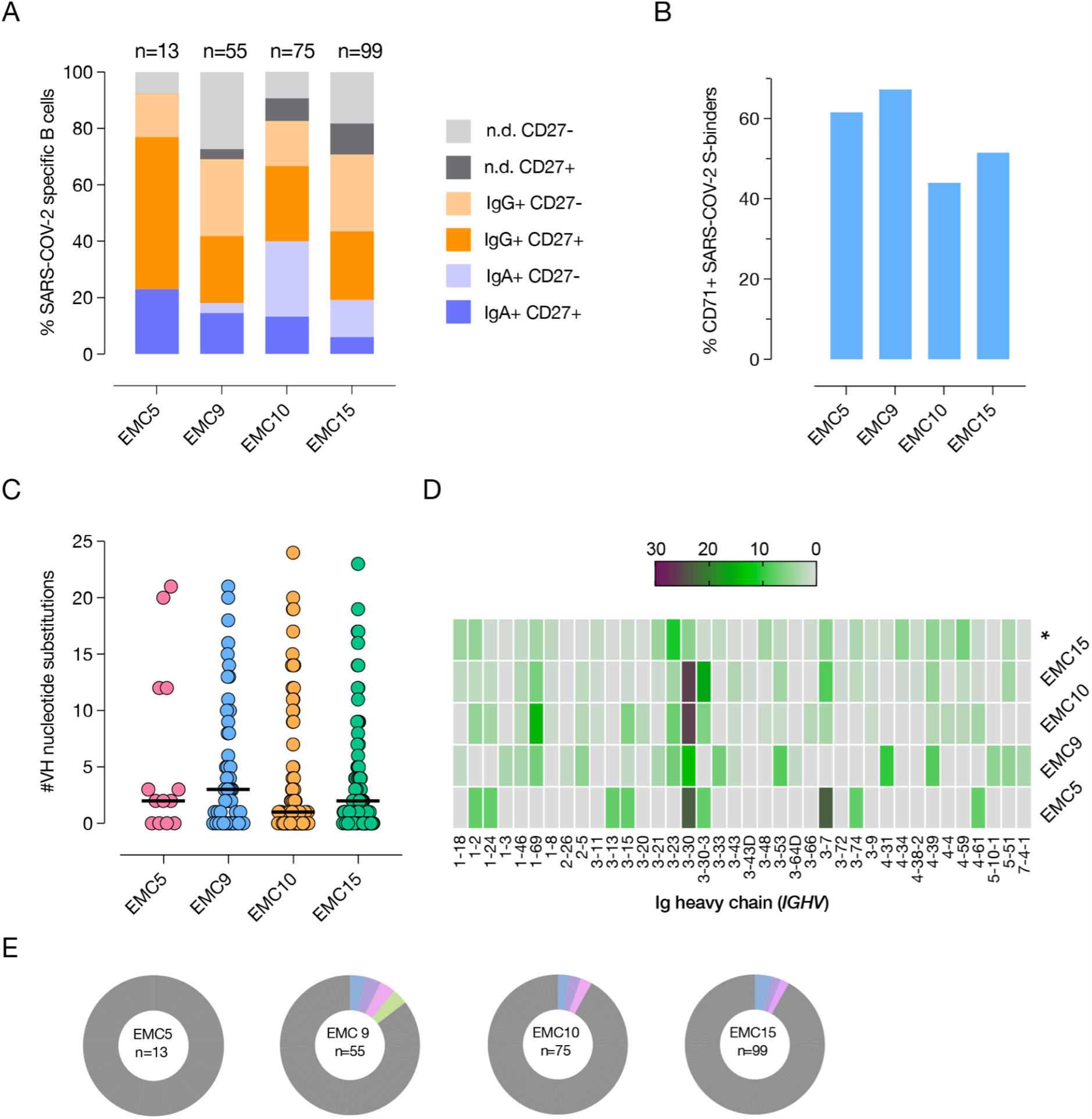
SARS-CoV-2 induces robust and diverse memory B-cell response in COVID-19 convalescent patients. Relative proportion of immunoglobulin isotype and classical memory B-cell surface marker CD27 expression (A) or activation marker CD71 expression (B) on B-cells from which SARS-CoV-2 S-specific mAbs were derived. n.d. = not determined. (C) Number of nucleotide substitutions in the VH genes of the mAbs isolated from the indicated donors. (D) VH germline gene (*IGHV*) usage in SARS-CoV-2 S-specific mAbs; * -indicates the frequency of germline gene usage in unselected human repertoire (37) for comparison. (E) Clonal lineage analysis of SARS-CoV-2 S-specific mAbs for each of the donors. Donor ID and the total number of clones are indicated in the center of each pie. Unique clones are shown in grey; colored slices indicate expanded clones.

**FIG S1.**
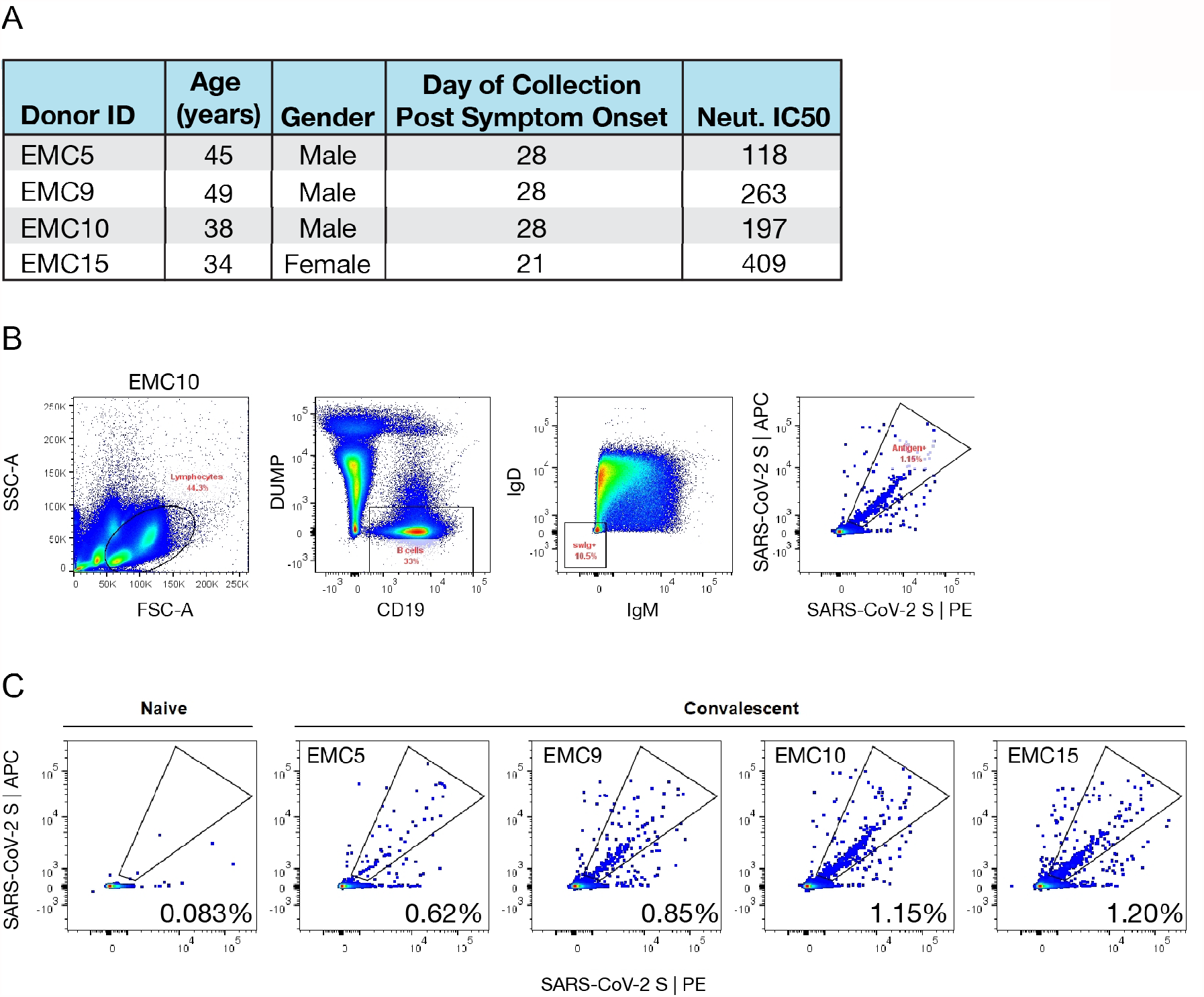
(A) Details of SARS-CoV-2 convalescent donors. Neutralization IC^50^ values are shown as reciprocal serum titers. (B-C) Identification and sorting of SARS-COV-2 S-reactive memory B-cells in COVID-19 convalescent patients. (B) Representative gating strategy for identification of class-switched SARS-CoV-2 S-reactive B-cells. (C) Percent of class-switched SARS-CoV-2 S-reactive B-cells in naïve and convalescent donors. The gated populations were single-cell sorted for cloning of the VH and VL genes. Naïve donor sample drawn and processed in August 2019 is shown for comparison.

### RBD-and NTD-targeting mAbs are potent neutralizers of SARS-CoV-2

From the cloned pairs of VH and VL genes of the four donors, we expressed and purified mAbs in a human IgG1 background using our yeast expression platform. These mAbs were screened for their S protein binding ability and the top 163 mAbs binders were evaluated for their neutralizing activity against authentic SARS-CoV-2 and rVSV-SARS2 at 100 nM and 10 nM antibody concentrations in micro-neutralization assays, respectively. A total of 44 mAbs neutralized authentic virus with more than 50% efficacy and 34 mAbs did the same for rVSV-SARS2 (**Fig. 2A**). Based on these data, we selected the top 40 nAbs for further analyses and used biolayer interferometry (BLI) to identify the specific domains of the S protein targeted by these nAbs. Most of the nAbs mapped to the RBD and included hACE2 competitors and non-competitors. However, we also identified multiple non-RBD binding nAbs including many that target the NTD (**Fig. 2B**). Finally, we ran 9-point neutralization curves on all 40 nAbs with authentic virus and rVSV-SARS2 to down-select to four each of the most potent RBD-and non-RBD targeting antibodies (data not shown). The neutralizing profiles of the RBD-targeting mAbs against rVSV-SARS2 and the authentic SARS-CoV-2 were very similar, with IC^50^ neutralization values in the picomolar range for ADI-56443 (75 pM) and ADI-56899 (89 pM) (**Fig. 2C**). As has been observed previously for NTD-targeting mAbs (25), rVSV-SARS2 neutralization curves for the non-RBD mAbs were shallower and these mAbs left an un-neutralized fraction of the virus (**Fig. 2D, right panel**). However, there was no un-neutralized fraction left with the authentic virus (**Fig. 2D, left panel**). ADI-56479 was the best NTD-targeting nAb with picomolar neutralization IC^50^ values against the authentic virus (144 pM).

**FIG 2.**
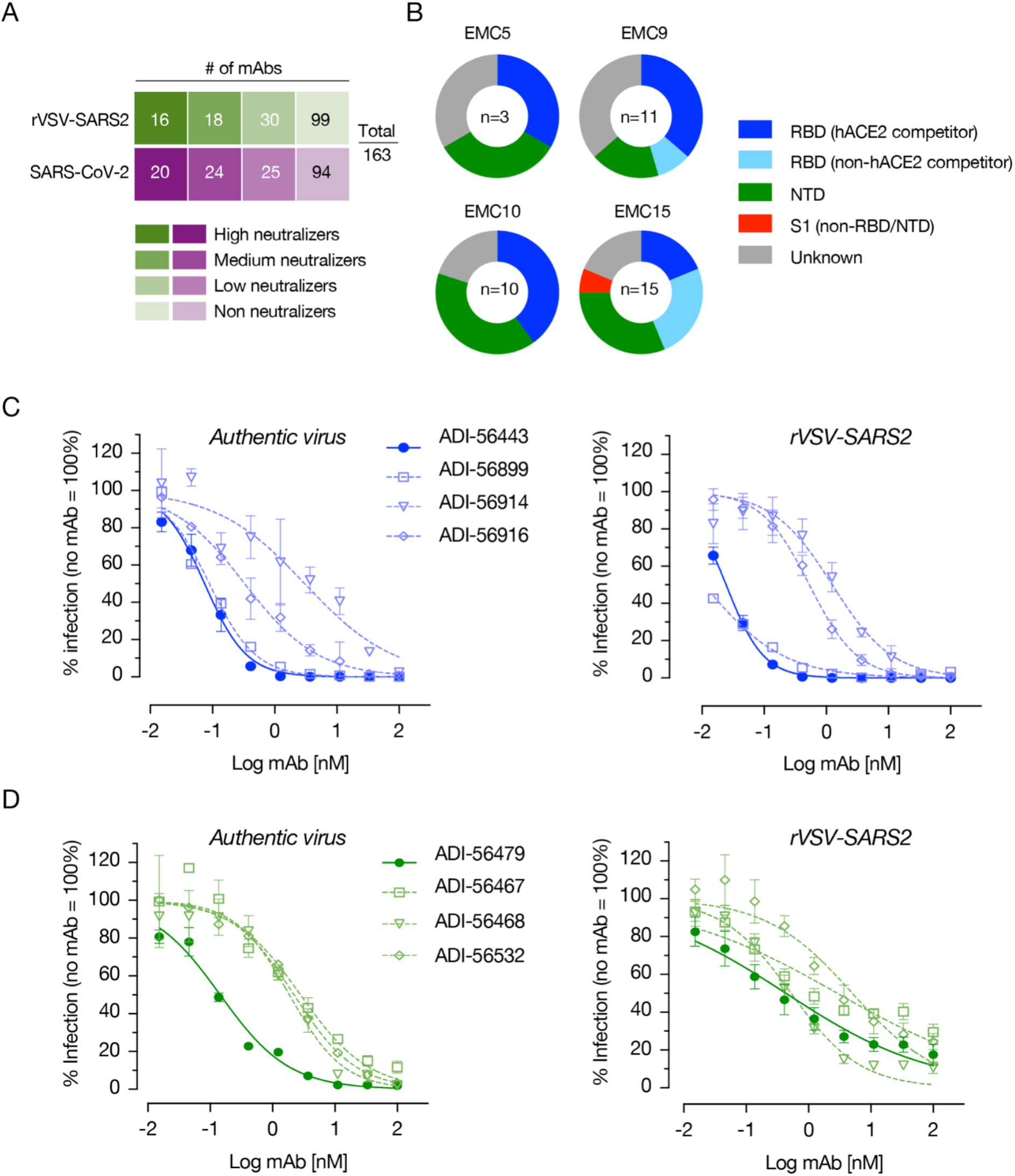
RBD-as well as NTD-targeting mAbs are potent neutralizers of SARS-CoV-2. (A)Screening of high affinity SARS-CoV-2 spike protein-binding mAbs for their neutralization capacity against rVSV-SARS2 and authentic SARS-CoV-2. The mAbs were divided into high neutralizers (80% neutralization efficacy), medium neutralizers (50-80%), low neutralizers (30-50%) and non-neutralizers (<30%) based on their capacity to neutralize rVSV-SARS2 at 10 nM or authentic virus at 100 nM antibody concentration. Number of mAbs in each group is indicated in the corresponding boxes. (B)Proportion of the 40 best nAbs, from each donor, targeting each of the indicated domains/regions of the spike protein. (C-D) Neutralization curves of each of the top four RBD-(C) and non-RBD-targeting (D) mAbs. The data were curve-fitted using a nonlinear regression (log [inhibitor] vs. normalized response, variable slope) to calculate IC^50^ values. (Average ± SD, n = 4 from two independent experiments for rVSV-SARS2 and n = 2 from a representative of multiple experiments for the authentic virus).

### Epitope mapping of the most potent RBD and NTD mAbs

To better understand the mechanism of neutralization by our most potent nAbs, we mapped their epitopes by selecting rVSV-SARS2 neutralization escape mutants. After serially passaging rVSV-SARS2 in the presence of increasing concentrations of the nAbs, we plaque-purified and sequenced resistant viruses to identify the S mutations that confer resistance. For the best RBD mAb (ADI-56443), a change at amino acid position 490 in the S protein (F490S) made rVSV-SARS2 highly resistant (>2,700-fold increase in neutralization IC^50^ value) to this mAb (**Fig. 3A**). To comprehensively map its epitope, we analyzed the binding capacity of ADI-56443 to a library of SARS-CoV-2 RBD single amino acid mutants displayed on the surface of yeast cells by flow cytometry. In addition to F490S, binding of ADI-56443 was completely abolished by C480S/R, E484K/G/D, C488Y/S and F490L/I/C RBD mutations (**Fig. 3B**). Mutations at other residues, including S494F in the RBD also significantly reduced this mAb’s binding (**Fig. 3B**). Remarkably, residues E484, F490, and S494 are shared with the epitope of mAb LY-CoV555 (Bamlanivimab), which received an EUA for COVID-19 treatment (38). E484K mutation is also present in multiple variants including P.1, P.2, B.1.525 and B.1.351 and viruses carrying this mutation are resistant to the currently used mAb therapy (39, 40). For the NTD mAb (ADI-56479), rVSV-SARS2 neutralization-escape mutations mapped to residues 145 (Y145D), 150 (K150E) and 152 (W152R) in the NTD and each one of these mutations individually afforded a ∼1,000 fold increase in the neutralization IC^50^ values (**Fig. 3C, left panel**). This was accompanied by a loss of binding of the ADI-56479 mAb to the mutant spike proteins as determined by ELISA using rVSV-SARS2 particles (**Fig. 3C, right panel**). Consistent with the RBD mAb (ADI-56443) being a competitor of hACE2-spike binding, its epitope partly overlaps with the receptor-binding interface (**Fig. 3D, right panel**). In contrast, all three of the residues in ADI-56479’s epitope are located in the N3 loop of NTD (Y145, K150 and W152) away from the RBD domain. Taken together, we have identified two potent neutralizing antibodies that target two distinct domains of the SARS-CoV-2 spike protein.

**FIG 3.**
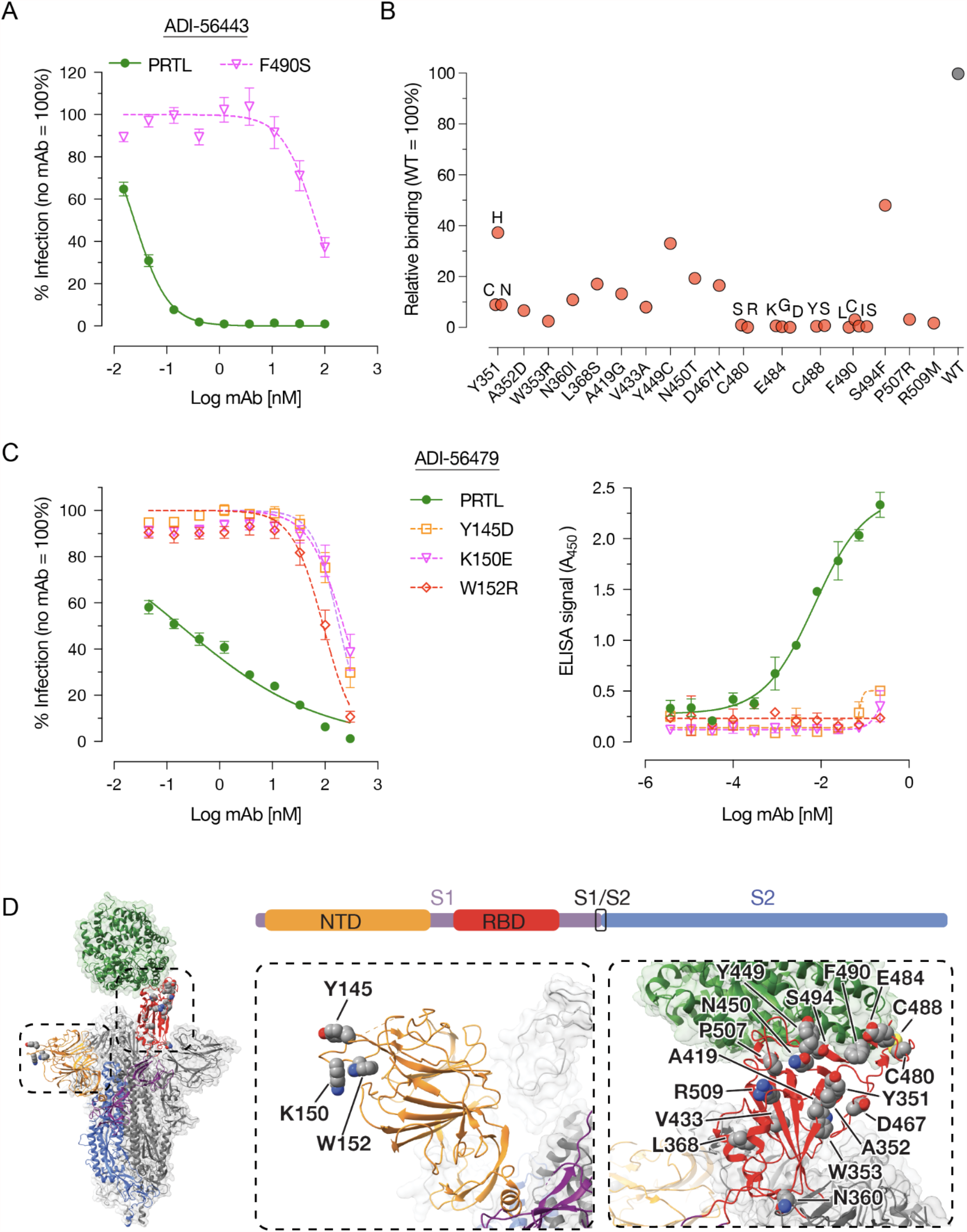
Epitope mapping of RBD-(ADI-56443) and NTD-targeting (ADI-56479) nAbs. (A) Pre-titrated amounts of the parental (PRTL) or indicated rVSV-SARS2 mutants were incubated with serial 3-fold dilutions of the ADI-56443 at room temperature for 1 h prior to infecting monolayers of Vero cells. After 7 h, cells were fixed, the nuclei were stained and infected cells were scored by eGFP expression. (Averages ± SEM, n = 4 from 2 independent experiments). (B) Binding capacity of ADI-56443 to a mutagenized library of SARS-CoV-2 RBD point mutants displayed on the surface of yeast cells was measured by flow cytometry. Key residues that led to a loss of ADI-56443 binding are shown (binding to cells displaying WT RBD was set at 100%). Antibody binding was assessed at their EC^80^ concentrations for the WT RBD construct. (C) Left panel -neutralization capacity of ADI-56473 against pre-titrated amounts of parental (PRTL) and indicated rVSV-SARS2 mutants was determined as above. (Averages ± SEM, n = 4 from 2 independent experiments). Right panel -ELISA plates coated with the parental or indicated rVSV-SARS2 mutants and binding of biotinylated-ADI-56749 to the spike protein was detected by using an HRP-conjugated streptavidin. A representative dataset from 2 independent experiments is shown here. (Average ± SD, n = 2). (D) Left panel: An overview of the SARS-CoV-2 S trimer bound to hACE2 (green) (26). For clarity, only the domains of one spike monomer have been colored (S1, purple; NTD, yellow; RBD, red; S2, Blue). Middle and right panels: a close-up view of the NTD (in yellow) and RBD (in red) with amino acid residues important for binding to ADI-56479 (middle panel) and ADI-56443 (right panel) are shown.

### rVSV-SARS2 NTD mutants are resistant to neutralization by COVID-19 convalescent sera

Although the most potent SARS-CoV-2 nAbs target the RBD (12, 13, 15–20), recent antibody profiling efforts and the emergence of multiple VOC with mutations in the NTD that affect the efficacy of nAbs suggesting that NTD-directed antibodies are important for effective control of virus infection (22–24, 41). Therefore, we tested if COVID-19 convalescent sera-mediated neutralization of rVSV-SARS2 is altered by the NTD mutations generated in response to the ADI-56479 mAb-driven selection (**Fig. 4**). Using a set of convalescent sera with high neutralizing activity (reciprocal serum neutralization IC^50^ titers of >500), we observed a significant effect of NTD point mutations on rVSV-SARS2 neutralization (**Fig. 4A**). Specifically, the sera showed a drop of 3.5-fold (Y145D) to 16-fold (K150E and W152R) in their neutralization IC^50^ titers as compared to the parental virus (**Fig. 4B**). Thus, our data further supports a significant role of the NTD-targeting antibodies in polyclonal sera-mediated SARS-CoV-2 neutralization (32, 33).

**FIG 4.**
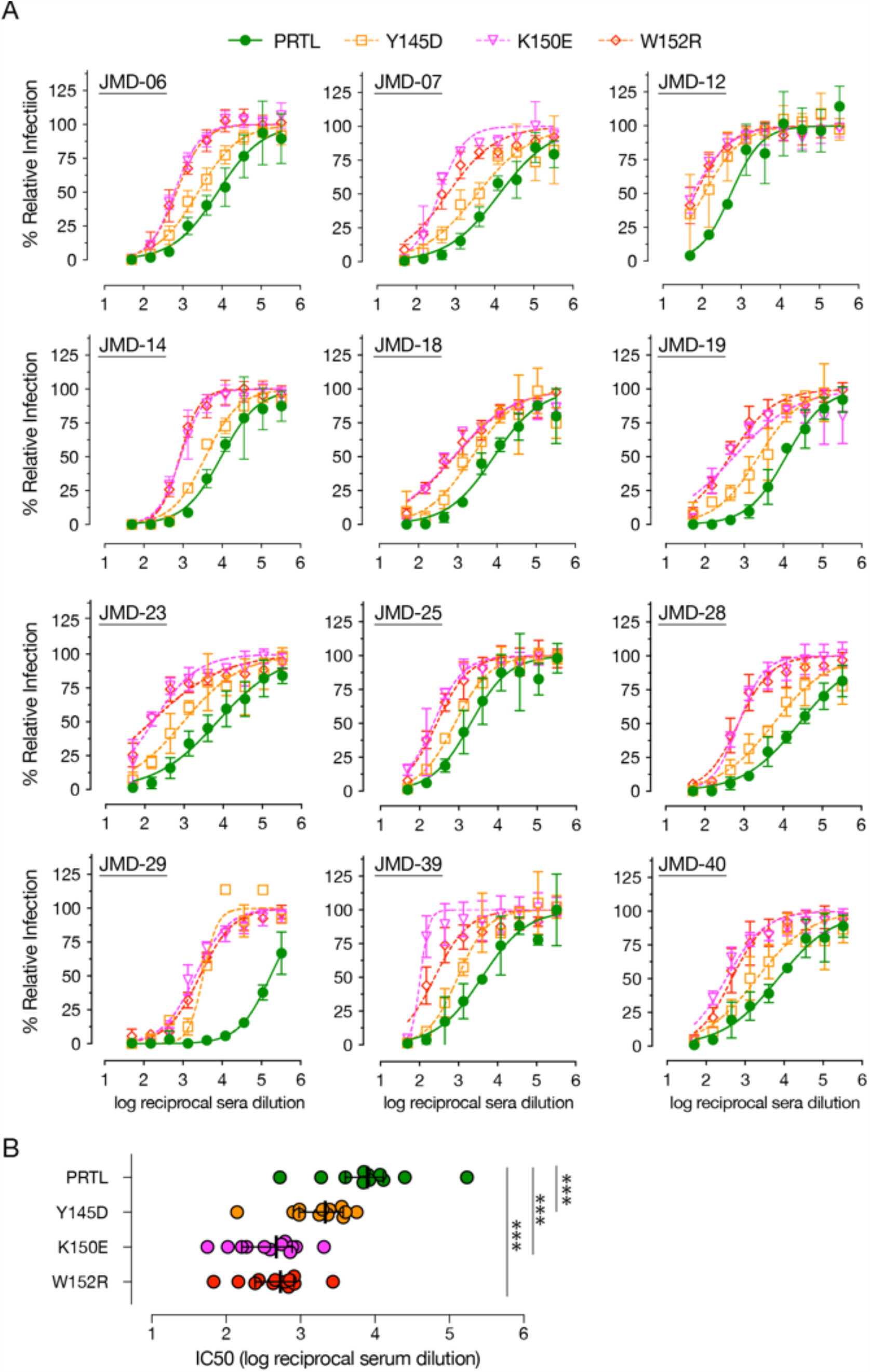
NTD point mutations significantly reduce neutralization of rVSV-SARS2 by convalescent sera. (A) Neutralization of the parental (PRTL) and indicated rVSV-SARS2 mutants with convalescent sera from 12 donors with high nAb antibody titers (reciprocal serum IC^50^ titers of >500). Pre-titrated amounts of indicated rVSV-SARS2 were incubated with 3-fold serial dilutions of COVID-19 convalescent sera at room temperature for 1 h and applied to monolayers of Vero cells. After 7 hours, cells were fixed and the nuclei were stained. Infected cells were scored by eGFP expression. (Averages ± SEM, n = 4 from 2 independent experiments). (B) Reciprocal serum neutralization IC50 titers of all the convalescent sera against parental (PRTL) and mutant rVSV-SARS2 shown in panel A are depicted. Wilcoxon test was performed to evaluate statistical significance between the neutralization efficacies against the parental and mutant viruses, *** -p≤ 0.001.

### Combination of RBD and NTD monoclonal antibodies limit the generation of neutralization-escape mutants

Since the top RBD (ADI-46443) and NTD (ADI-56479) nAbs bind to two distinct domains of the S protein (**Fig. 3D**), we reasoned that combining them may enhance their neutralization efficacy. Given the vastly different IC^50^ values of the two mAbs with rVSV-SARS2, we were not able to calculate a classical synergy combination index (CI) analysis based on their equimolar combination at IC^50 (42)^. However, we did observe a modest left shift in the rVSV-SARS2 neutralization curve by combining each mAb starting at their IC^90^ concentrations (**Fig. 5A**). Next, we evaluated if combining the two mAbs, in 1:1 ratio at their neutralization IC^90^ concentrations, also has an effect on limiting the emergence of rVSV-SARS2 neutralization-escape mutants as compared to the single mAbs (**Fig. 5B**). As expected, rVSV-SARS2 rather easily escaped individual mAbs reaching a peak titer of 3×10^6^ (RBD nAb) and 2.7×10^5^ (NTD nAb) infectious units per mL as compared to the no nAb controls (1.2×10^7^ infectious units per mL) at 48 h post-infection (**Fig. 5C**). However, rVSV-SARS2 failed to grow in the presence of the mAb combination (peak titer of 72 infectious units per mL). Thus, in addition to enhancing neutralization efficacy, this mAb combination limits the development of mAb-resistant spike mutants.

**FIG 5.**
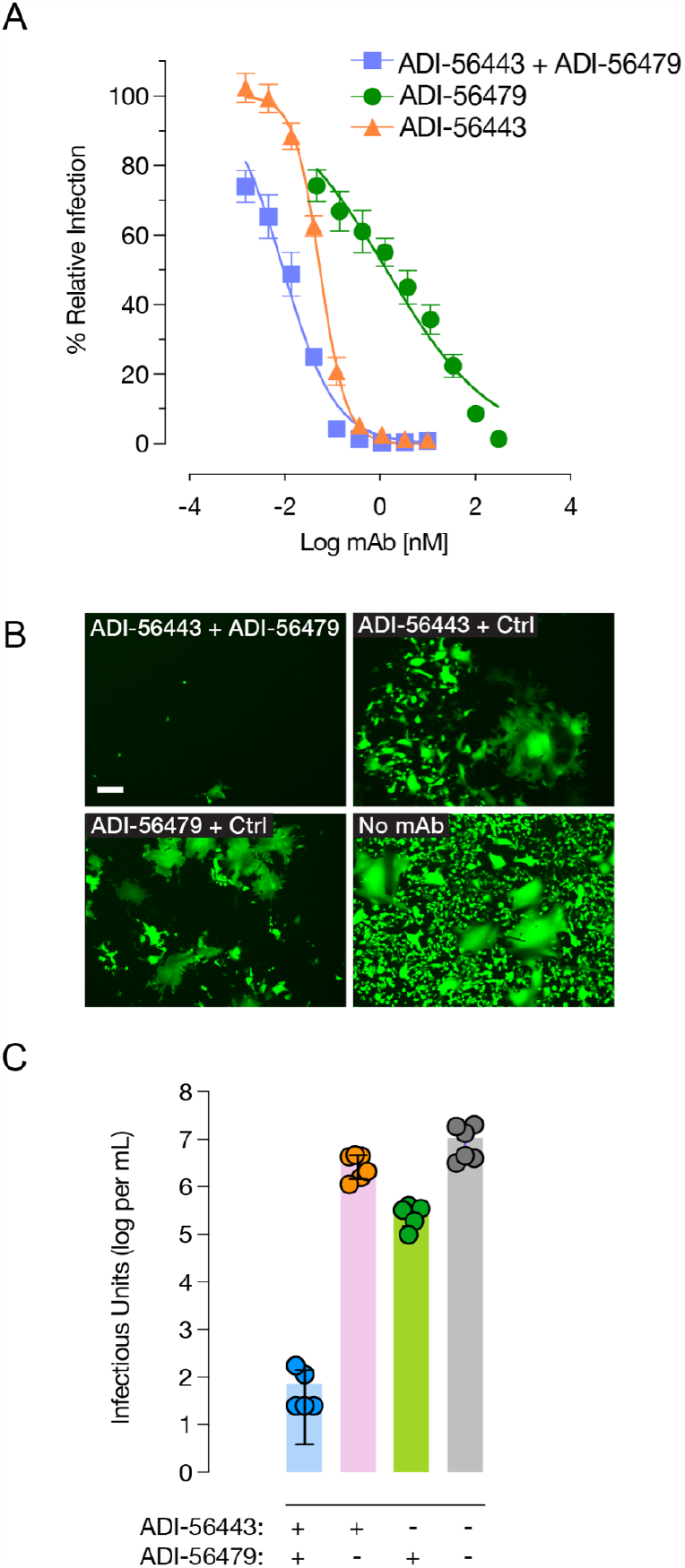
Combining RBD-and NTD-targeting mAbs enhances resistance to neutralization-escape. (A) Neutralization of rVSV-SARS2 by RBD-(ADI-56443) and NTD-(ADI-56479) targeting mAbs used alone or in combination. (Average ± SD, n = 5-6 from two independent experiments). (B) Representative images of eGFP expression of Vero cells at 24 hpi with rVSV-SARS2 in the presence of IC^90^ concentrations of the RBD-(ADI-56443) and NTD-(ADI-56479) targeting mAbs alone or in combination. No mAb control is shown for comparison. Scale bar = 100 µm (C) Quantitation of rVSV-SARS2 (infectious units per mL) produced from Vero cells in panel B at 48 hpi. (Average ± SD, n = 5-6 from two independent experiments).

## Discussion

All of the SARS-CoV-2 nAbs in phase III clinical trials or clinical use under EUA to treat COVID-19 are RBD-specific (12, 13, 15–18, 20). However, potent non-RBD nAbs are present in COVID-19–convalescent patients, and the emergence of NTD mutations that afford resistance to neutralization in multiple SARS-CoV-2 VOC highlights the importance of NTD as an important target for nAbs. Here, we isolated potent RBD and NTD nAbs from multiple COVID-19 donors and mapped their epitopes in the S protein. We show that neutralization-escape mutations to the NTD nAb significantly reduce the efficacy of polyclonal COVID-19 convalescent sera. Importantly, a combination of RBD and NTD mAbs modestly increased efficacy and could effectively limit the emergence of nAb-resistant S mutations.

Consistent with the increasingly recognized importance and prevalence of non-RBD nAbs (41), 12 of our top 40 nAbs targeted NTD (**Fig. 1 and 2**). Six of these NTD nAbs were derived from two variable heavy (VH) genes (VH1-24 or VH3-30), suggesting their preponderance in NTD nAbs. Some NTD nAbs require significant somatic hypermutations for their efficacy (23, 27). However, 11 out of the 12 nAbs that we discovered carried ≤3 somatic hypermutations, indicating that they arose at an early time-point post-infection. Thus, the nAb response to NTD is polyclonal and many potent NTD nAbs require relatively low levels of affinity maturation, as described recently (23).

Although blocking of hACE2:Spike interactions is one of the major mechanisms of virus neutralization, NTD-targeting antibodies with other blocking mechanisms have been reported (23–28). However, their precise mechanisms of action remain unclear. Some appear to inhibit a post-attachment phase of virus infection and may block subsequent viral steps in entry and/or promote Fc-mediated effector functions (23, 24). For the distantly related MERS-CoV, the NTD can bear critical epitope(s) for neutralization (43). Here, we show that a potent NTD nAb (ADI-56479) interacts with residues Y145, K150 and W152 in the NTD (**Fig. 3**), which are located in the N3 loop of an ‘antigenic supersite.’ Other potent NTD nAbs also recognize the same antigenic supersite (17, 23–25, 27). Recently, Suryadevara et al. (24) showed that mutations F140S, and G142D or R158S in the NTD confer resistance of two other NTD nAbs. Notably, our rVSV-SARS2 neutralization-escape variants (Y145D, K150E and W152R) were significantly resistant to neutralization by convalescent COVID-19 sera (**Fig. 4**). In addition, mutations at N148S, K150R/E and S151P in the NTD epitope exhibited reductions in sensitivity to three COVID-19 convalescent plasma samples (32). A deletion at F140 that occurred in response to immune pressure from highly neutralizing convalescent plasma also partially reduced neutralization (33). Interestingly, a VOC, B.1.429, which displays moderate reduction in neutralization against convalescent and post-vaccination sera carries a mutation at W152 (44). One of the Indian variants, B.1.617.2, has as well shown deletions at position 156 and 157 emphasizing that the NTD is acquiring adaptive mutations that counteract the immune response (22, 44). Collectively, these findings underscore the importance of NTD in virus entry and the potential of NTD mutations to impact the effectiveness of vaccines and nAb therapies.

Combination therapies with RBD and NTD nAbs have been proposed as a strategy to mitigate the emergence of antibody-resistant variants (6, 17, 24, 25, 32). We observed that a combination of RBD and NTD mAbs enhanced neutralization efficacy relative to the individual mAbs. In particular, the virus was less likely to escape when passaged in the presence of a RBD and NTD nAb cocktail than in the presence of each nAb alone (**Fig. 5**). This finding is consistent with what has recently been reported with other NTD and RBD nAb combinations (23, 24). Our findings, together with recent publications, provide evidence that some NTD-targeting mAbs can efficiently inhibit SARS-CoV-2 infection. While SARS-CoV-2 continues to evolve into multiple VOC, a combination antibody therapy with RBD and NTD nAbs could provide an important solution for treating COVID-19 and impede the generation of novel variants especially in populations that cannot be vaccinated.

## Material and Methods

### Cells and viruses

The African vervet monkey kidney Vero (CCL-81) cells were cultured in Dulbecco’s Modified Eagle Medium (DMEM high glucose, Gibco) supplemented with 2% heat-inactivated fetal bovine serum (FBS, Atlanta Biologicals), 1% Penicillin/Streptomycin (Gibco) and 1% Gluta-MAX (Gibco). The cells were passaged every three to four days using 0.05% Trypsin/EDTA solution (Gibco).

All the experiments described here with the authentic SARS-CoV-2 (Washington state isolate MT020880.1) were carried out in BSL-3 laboratories at USAMRIID, Frederick, MD as per Federal regulations under institutional biosafety committee-approved protocols. Virus stocks were prepared as described previously (31). Sequencing of the virus stock revealed a single mutation (a histidine to tyrosine change at amino acid position 655, H655Y) in the spike glycoprotein relative to the reference Washington state isolate.

A plaque-purified rVSV-SARS2 corresponding to the passage 9 (plaque #2 virus) described previously (31) was used for these studies. It is referred to here as the parental (PRTL) virus. Virus stocks were generated by growing the virus on Vero cells. Appropriate approvals from the Environmental Health and Safety Department and the Institutional Biosafety Committee at Albert Einstein College of Medicine were sought for using rVSV-SARS2 at biosafety level 2.

### Collection of COVID-19 convalescent donor blood samples

Convalescent blood samples were collected from healthy adult patient volunteers who had mild COVID-19 and a positive RT-qPCR test for SARS-CoV-2 in March 2020 in Westchester County, New York. These patients were neither hospitalized nor required oxygen supplementation during illness. All donors recovered and were asymptomatic for at least 14 days prior to venipuncture to collect serum and PBMCs. Serum was centrifuged, aliquoted and stored at -80°C. Sera were heat-inactivated at 56°C for 30 min and stored at 4°C prior to antibody testing. The study protocol was approved by the Institutional Review Board (IRB) of the Albert Einstein College of Medicine (IRB number 2016-6137).

### RT-qPCR to detect SARS-CoV-2 infection

SARS-CoV-2 RT-qPCR was performed as per the CDC protocols (45). Briefly, RNA was isolated from blood and PBMCs using Quick-RNA Viral Kit (Zymo). Total RNA was mixed with the respective primers and probes (all purchased from IDT) specific for 2019-nCoV (N1 and N2 assays), SARS-like coronaviruses (N3 assay), and human RNase P (RP assay) together with TaqPath™ 1-Step RT-qPCR Master Mix CG (ThermoFisher). A plasmid containing the complete nucleocapsid gene from 2019-nCoV (IDT) was used as a positive control. In addition, RNA transcribed from a plasmid containing a portion of the RPP30 gene (IDT) was used as quality control for the RNA isolation. All samples were run and analyzed using the iQ™5 device (BioRad).

### rVSV-SARS2 neutralization assay

Parental and mutant rVSV-SARS2 were generated and used for microneutralization assay as described previously (31). In short, serum samples or monoclonal antibodies were serially diluted and incubated with virus for 1 h at room temperature. For initial screening, a single concentration of 10 nM mAb was used instead. Serum or antibody-virus mixtures were then added in duplicates or triplicates to 96-well plates (Corning) containing monolayers of Vero cells. After 7 h at 37°C and 5% CO2, cells were fixed with 4% paraformaldehyde (Sigma), washed with PBS, and stored in PBS containing Hoechst-33342 (Invitrogen). Viral infectivity was measured by automated enumeration of GFP-positive cells from captured images using a Cytation5 automated fluorescence microscope (BioTek) and analyzed using the Gen5 data analysis software (BioTek). The half-maximal inhibitory concentration (IC50) of the mAbs or sera was calculated using a nonlinear regression analysis with GraphPad Prism software.

For the mAb combination experiment, NTD (300 nM) or RBD (10 nM) mAbs were combined with each other or with an irrelevant mAb at equivalent concentration. Three-fold serial dilutions of the mAbs were serial dilutions of these mixtures were then tested for their neutralization capacity.

### SARS-CoV-2 neutralization assay

Neutralization assay using authentic virus was performed as described previously (31). In brief, mAbs with an initial concentration of 100 nM were serially diluted, mixed with pre-titrated amounts of SARS-CoV-2 (MOI = 0.2) and incubated for 1 h at 37°C and 5% CO2. The inoculum was added to Vero-E6 cell monolayers in 96 well plates and incubated for 1 h at 37°C and 5% CO2. For initial screening, a single concentration of 100 nM was used instead. The virus:serum inoculum was removed, cells were washed with PBS and media was added. At 24 hours post-infection, cells were treated with 10% paraformaldehyde, washed with PBS and permeabilized with 0.2% Triton-X for 10 min at room temperature. Cells were immunostained with SARS-CoV-1 nucleocapsid protein-specific antibody (Sino Biologic; Cat# 40143-R001) and AlexaFluor 488 labeled secondary antibody. Stained cells were imaged using an Operetta (Perkin Elmer) high content imaging instrument and the number of infected cells were determined using Harmony Software (Perkin Elmer).

### Isolation of PBMCs

Approximately 64 mL of whole blood collected from each donor using 8 mL BD Vacutainer CPT sodium heparin mononuclear cell preparation tubes was stored upright at room temperature for >1 h prior to centrifugation. Samples were centrifuged using a swinging bucket centrifuge at room temperature for 30 minutes at 4°C at 1800 RCF. The mononuclear cell layer was removed by using a pipette and pooled together in a single tube for each donor. The total volume was brought to 45 mL with Mg2+ and Ca2+ free Hank’s Balanced Salt Solution (HBSS). Cells were resuspended by inverting the tubes and centrifuged at room temperature for 10 minutes at 330 RCF. After removing the supernatant, the cells were resuspended in 90% FBS supplemented with 10% DMSO to a final concentration of 1×10^7^ cells per mL and stored at -150°C.

### Human ACE2 and SARS-CoV-2 spike antigens

Prefusion-stabilized SARS-CoV-2 S-2P spike ectodomain (residues 1-1208) was expressed and purified as described previously (7). Plasmids encoding residues 1-305 of the SARS-CoV-2 spike with a C-terminal HRV3C cleavage site, monomeric Fc-tag and 8x HisTag (SARS-CoV-2 NTD); residues 319-591 of the SARS-CoV-2 spike with a C-terminal HRV3C cleavage site, monomeric Fc-tag and 8x HisTag (SARS-CoV-2 RBD-SD1); residues 1-615 of human ACE2 with a C-terminal 8x HisTag and TwinStrepTag (hACE2) were transfected into FreeStyle-293F cells. Cell supernatants were harvested after 6 days and expressed proteins were purified by affinity chromatography using a Superdex 200 Increase column (Cytiva). The SARS-CoV-2 NTD and RBD-SD1 proteins were purified using Protein A resin (Pierce), whereas hACE2 protein was purified using StrepTactin resin (IBA). These proteins were then further purified by size-exclusion chromatography using a buffer composed of 2 mM Tris pH 8.0, 200 mM NaCl and 0.02% NaN3. The SARS-CoV-2 S1 subunit (Cat# S1N-C52H3) was purchased from Acro Biosystems.

### Sorting of SARS-CoV-2 spike-reactive single B-cells

B-cells were purified and sorted as described previously (19). In short, B-cells purified from donor PBMCs using the MACS Human B-Cells isolation kit (Miltenyi Biotec Miltenyi Biotec Cat# 130-091-151) were stained with a panel of antibodies: anti-human CD19 (PE-Cy7; Biolegend Cat# 302216), CD3 (PerCP-Cy5.5; Biolegend Cat# 30040), CD8 (PerCP-Cy5.5; Biolegend Cat# 344710), CD14 (PerCP-Cy5.5; Invitrogen Cat# 45-0149-42), CD16 (PerCP-Cy5.5; Biolegend Cat# 360712), IgM (BV711; BD

Biosciences Cat# 747877), IgD (BV421; Biolegend Cat# 348226), IgA (AF-488; Abcam Cat# Ab98553), IgG (BV605; BD Biosciences Cat#563246), CD27 (BV510; BD Biosciences Cat# 740167), CD71 (APC-Cy7; Biolegend Cat# 334110), propidium iodide (PI), and an equimolar mixture of APC-and PE-labeled SARS-CoV-2 S-2P protein tetramers. BD FACS Aria II Fusion (BD Biosciences) was used for index sorting. The class-switched B-cells were defined as CD19^+^CD3^−^CD8^−^CD14^−^CD16^−^PI^−^IgM^−^IgD^−^ cells with a reactivity to APC-and PE-labeled SARS-CoV-2 S-2P tetramers. Single cells were sorted and plates were stored at -80°C till further processing. Flow cytometry data was analyzed using FlowJo software.

### Amplification and cloning of IgG variable heavy and light chain genes

Human antibody variable gene transcripts (VH, Vκ, Vλ) were amplified and cloned as previously described (34). Briefly, reverse transcription polymerase chain reaction (RT-PCR) (SuperScript IV enzyme (Thermo Scientific) followed by nested PCR (HotStarTaq Plus DNA Polymerase (Qiagen) with cocktails of variable region and IgM-, IgD-, IgA-and IgG-specific constant-region primers was performed. The next nested PCR was carried out to allow cloning by homologous recombination and amplified gene transcripts were transformed into *S. cerevisiae* (46). Finally, yeast cells were washed with sterile water, resuspended in selective media and plated. The individual yeast clones were analyzed using Sanger sequencing.

### Expression and purification of human mAbs

mAbs were expressed as full-length human IgG1 proteins in *S. cerevisiae* cultures, as previously described (34). Briefly, yeast cultures were grown in a 24-well format at 30°C and 80% relative humidity with shaking at 650 RPM in Infors Multitron shakers. Culture supernatants were harvested after 6 days of growth and IgGs were purified by protein A-affinity chromatography. IgGs bound to the agarose were eluted with 200 mM acetic acid with 50 mM NaCl (pH 3.5) and neutralized with 1/8 (v/v) 2 M HEPES (pH 8.0).

### Biolayer interferometry (BLI) to assess mAb:antigen binding

As previously described (34), for apparent equilibrium dissociation constant (KDApp) determination ForteBio Octet HTX instrument (Molecular Devices(34) was used to measure the biolayer interferometry (BLI) kinetic of IgG binding to recombinant antigens. In short, the IgGs were captured on anti-human IgG (AHC) biosensors (Molecular Devices). For BLI measurements involving Strep-tagged antigens, the sensors were additionally incubated in a biocytin solution to saturate remaining streptavidin binding sites. After a one-minute baseline step, IgG-loaded biosensors were exposed for 180 secs to the recombinant antigen. Next, the dissociation of the antigen from the biosensor surface was measured. For binding responses >0.1 nm, data were aligned, inter-step corrected to the association step, and fit to a 1:1 binding model using the ForteBio Data Analysis Software, version 11.1.

### mAb competition assay using BLI

Competitive binding of mAbs to recombinant SARS-CoV-2 RBD with hACE2 was evaluated using the ForteBio Octet HTX instrument (Molecular Devices) as described previously (34). Briefly, IgGs were captured onto AHC biosensors (Molecular Devices) to achieve a sensor response between 1-1.4 nm followed by an inert IgG to occupy any remaining binding sites on the biosensor. The sensors were then equilibrated for a minimum of 30 min. The loaded sensors were additionally exposed to hACE2 for 90 secs, prior to the binning analysis to assess any interactions between secondary molecules and proteins on the sensor surface. After a 60-seconds baseline step, association to recombinant SARS-CoV-2 RBD was performed for 180 secs and was finally followed by exposure to hACE2. The data was analyzed using the ForteBio Data Analysis Software version 11.0. The absence of binding by the secondary molecule indicates an occupied epitope (competitor) and binding indicates a non-competing antibody.

### Epitope mapping using a yeast-display library

Epitope mapping was done using a library of SARS-CoV-2 RBD point mutants displayed on the yeast surface as described previously (47). To select for mutants in the RBD-SD1 library with diminished binding to ADI-56443, the mutant RBD-SD1 library and WT RBD-SD1 yeast were incubated with ADI-56443 at its EC^80^ concentration. Yeast cells from the library expressing HA-tag but with reduced ADI-56443 binding, as compared to WT RBD SD1, were sorted by using a BD FACS Aria II (Becton Dickerson). The sorted cells were propagated and the selection process was repeated to further enrich yeast cells encoding RBD mutants with reduced ADI-56443-binding. RBD sequences in the cell clones were sequenced and those possessing single amino acid substitutions were cultured, protein-expression induced, and evaluated for their binding to ADI-56643 at EC^80^ concentration by flow cytometry. Binding signal was normalized to that of the reference WT RBD-SD1 (set as 100%).

### Selection of neutralization-escape mutants of rVSV-SARS2

Generation of rVSV-SARS2 that escaped neutralization with our top RBD (ADI-56443) and NTD (ADI-56479) mAbs was done as described previously (19). Briefly, rVSV-SARS2 was pre-incubated with IC^90^ concentrations of ADI-56443 (0.37 nM) and ADI-56479 (100 nM) were applied to monolayers of Vero cells and infection was allowed to proceed in the presence of the mAbs. Virus supernatants were harvested from the cells at 48-72 hpi and passaged again by doubling the amount of antibody for subsequent passage. After 3 passages, mAb-resistant viruses were plaque-purified, their phenotypes were confirmed by neutralization assay and S gene sequences were determined by RT-PCR followed by Sanger sequencing as described previously (31).

### Assay for resistance to the generation of neutralization-escape rVSV-SARS2 mutants

rVSV-SARS2 particles (MOI = 0.05) were incubated with IC^90^ concentrations of ADI-56443 (0.37 nM) and ADI-56479 (100 nM) for 1 hr at room temperature and the virus-antibody mixture was used to infect Vero cells in 6-well plates. Cells were imaged for eGFP expression at 24 hpi. Virus supernatants were harvested at 48 hpi and the amount of virus produced was determined by titration on Vero cells in the absence of mAbs as described previously (31).

### ELISA to detect binding of NTD mAbs to rVSV-SARS2 particles

High-protein binding 96-well ELISA plates (Corning) were coated with normalized amounts of purified parental or the mutant rVSV-SARS2 overnight at 4°C, and blocked with 3% nonfat dry milk in PBS (PBS-milk) for 1 h at 37°C. Plates were washed and incubated with biotinylated ADI-56479 at a concentration starting 0.22 ug/mL with serial 3-fold dilutions in 1% PBS-milk supplemented with 0.1% Tween-20 for 1 h at 37°C. Plates were washed three times and incubated with streptavidin-HRP (Pierce Cat#21130) diluted 1:3000 in 1% PBS-milk supplemented with 0.1% Tween-20 for 1 h at 37C and detected using 1-Step Ultra TMB-ELISA Substrate Solution (Thermo Fisher Scientific). Plates were read using a Cytation 5 imager (BioTek) at 450 nm.

## Acknowledgements

We thank I. Gutierrez, E. Valencia, and L. Polanco for laboratory management. This work was supported in part by National Institutes of Health (NIH) grants R01AI132633 (to K.C.), R01AI125462 (to J.R.L.) and R21AI141367 (to J.P.D). M.E.D. was a Latin American Fellow in the Biomedical Sciences, supported by the Pew Charitable Trusts. R.H.B.III. and R.J.M. were partially supported by the NIH training grant 2T32GM007288-45 (Medical Scientist Training Program) at Albert Einstein College of Medicine. K.C. and J.R.L. were also supported by an Einstein Pilot Project grant for SARS-CoV2.

## Bibliography

1. Wu F, Zhao S, Yu B, Chen Y-M, Wang W, Song Z-G, Hu Y, Tao Z-W, Tian J-H, Pei Y-Y, Yuan M-L, Zhang Y-L, Dai F-H, Liu Y, Wang Q-M, Zheng J-J, Xu L, Holmes EC, Zhang Y-Z. 2020. A new coronavirus associated with human respiratory disease in China. Nature 579:265–269.

2. Dong E, Du H, Gardner L. 2020. An interactive web-based dashboard to track COVID-19 in real time. Lancet Infect Dis 20:533–534.

3. Siemieniuk RA, Bartoszko JJ, Ge L, Zeraatkar D, Izcovich A, Kum E, Pardo-Hernandez H, Rochwerg B, Lamontagne F, Han MA, Liu Q, Agarwal A, Agoritsas T, Chu DK, Couban R, Darzi A, Devji T, Fang B, Fang C, Flottorp SA, Cusano E. 2020. Drug treatments for covid-19: living systematic review and network meta-analysis. BMJ 370:m2980.

4. Challen R, Brooks-Pollock E, Read JM, Dyson L, Tsaneva-Atanasova K, Danon L. 2021. Risk of mortality in patients infected with SARS-CoV-2 variant of concern 202012/1: matched cohort study. BMJ 372:579.

5. Li F. 2016. Structure, function, and evolution of coronavirus spike proteins. Annu Rev Virol 3:237–261.

6. Walls AC, Park Y-J, Tortorici MA, Wall A, McGuire AT, Veesler D. 2020. Structure, Function, and Antigenicity of the SARS-CoV-2 Spike Glycoprotein. Cell 181:281–292.e6.

7. Wrapp D, Wang N, Corbett KS, Goldsmith JA, Hsieh C-L, Abiona O, Graham BS, McLellan JS. 2020. Cryo-EM structure of the 2019-nCoV spike in the prefusion conformation. Science 367:1260–1263.

8. Krempl C, Schultze B, Laude H, Herrler G. 1997. Point mutations in the S protein connect the sialic acid binding activity with the enteropathogenicity of transmissible gastroenteritis coronavirus. J Virol 71:3285–3287.

9. Künkel F, Herrler G. 1993. Structural and functional analysis of the surface protein of human coronavirus OC43. Virology 195:195–202.

10. Lu G, Wang Q, Gao GF. 2015. Bat-to-human: spike features determining “host jump” of coronaviruses SARS-CoV, MERS-CoV, and beyond. Trends Microbiol 23:468–478.

11. Barnes CO, West AP, Huey-Tubman KE, Hoffmann MAG, Sharaf NG, Hoffman PR, Koranda N, Gristick HB, Gaebler C, Muecksch F, Lorenzi JCC, Finkin S, Hägglöf T, Hurley A, Millard KG, Weisblum Y, Schmidt F, Hatziioannou T, Bieniasz PD, Caskey M, Bjorkman PJ. 2020. Structures of Human Antibodies Bound to SARS-CoV-2 Spike Reveal Common Epitopes and Recurrent Features of Antibodies. Cell 182:828–842.e16.

12. Baum A, Ajithdoss D, Copin R, Zhou A, Lanza K, Negron N, Ni M, Wei Y, Mohammadi K, Musser B, Atwal GS, Oyejide A, Goez-Gazi Y, Dutton J, Clemmons E, Staples HM, Bartley C, Klaffke B, Alfson K, Gazi M, Kyratsous CA. 2020. REGN-COV2 antibodies prevent and treat SARS-CoV-2 infection in rhesus macaques and hamsters. Science 370:1110–1115.

13. Hansen J, Baum A, Pascal KE, Russo V, Giordano S, Wloga E, Fulton BO, Yan Y, Koon K, Patel K, Chung KM, Hermann A, Ullman E, Cruz J, Rafique A, Huang T, Fairhurst J, Libertiny C, Malbec M, Lee W-Y, Kyratsous CA. 2020. Studies in humanized mice and convalescent humans yield a SARS-CoV-2 antibody cocktail. Science 369:1010–1014.

14. Hassan AO, Case JB, Winkler ES, Thackray LB, Kafai NM, Bailey AL, McCune BT, Fox JM, Chen RE, Alsoussi WB, Turner JS, Schmitz AJ, Lei T, Shrihari S, Keeler SP, Fremont DH, Greco S, McCray PB, Perlman S, Holtzman MJ, Diamond MS. 2020. A SARS-CoV-2 Infection Model in Mice Demonstrates Protection by Neutralizing Antibodies. Cell 182:744–753.e4.

15. Cao Y, Su B, Guo X, Sun W, Deng Y, Bao L, Zhu Q, Zhang X, Zheng Y, Geng C, Chai X, He R, Li X, Lv Q, Zhu H, Deng W, Xu Y, Wang Y, Qiao L, Tan Y, Xie XS. 2020. Potent Neutralizing Antibodies against SARS-CoV-2 Identified by High-Throughput Single-Cell Sequencing of Convalescent Patients’ B Cells. Cell 182:73–84.e16.

16. Ju B, Zhang Q, Ge J, Wang R, Sun J, Ge X, Yu J, Shan S, Zhou B, Song S, Tang X, Yu J, Lan J, Yuan J, Wang H, Zhao J, Zhang S, Wang Y, Shi X, Liu L, Zhang L. 2020. Human neutralizing antibodies elicited by SARS-CoV-2 infection. Nature 584:115–119.

17. Liu L, Wang P, Nair MS, Yu J, Rapp M, Wang Q, Luo Y, Chan JF-W, Sahi V, Figueroa A, Guo XV, Cerutti G, Bimela J, Gorman J, Zhou T, Chen Z, Yuen K-Y, Kwong PD, Sodroski JG, Yin MT, Ho DD. 2020. Potent neutralizing antibodies against multiple epitopes on SARS-CoV-2 spike. Nature 584:450– 456.

18. Pinto D, Park Y-J, Beltramello M, Walls AC, Tortorici MA, Bianchi S, Jaconi S, Culap K, Zatta F, De Marco A, Peter A, Guarino B, Spreafico R, Cameroni E, Case JB, Chen RE, Havenar-Daughton C, Snell G, Telenti A, Virgin HW, Corti D. 2020. Cross-neutralization of SARS-CoV-2 by a human monoclonal SARS-CoV antibody. Nature 583:290–295.

19. Wec AZ, Wrapp D, Herbert AS, Maurer DP, Haslwanter D, Sakharkar M, Jangra RK, Dieterle ME, Lilov A, Huang D, Tse LV, Johnson NV, Hsieh C-L, Wang N, Nett JH, Champney E, Burnina I, Brown M, Lin S, Sinclair M, Walker LM. 2020. Broad neutralization of SARS-related viruses by human monoclonal antibodies. Science.

20. Zost SJ, Gilchuk P, Chen RE, Case JB, Reidy JX, Trivette A, Nargi RS, Sutton RE, Suryadevara N, Chen EC, Binshtein E, Shrihari S, Ostrowski M, Chu HY, Didier JE, MacRenaris KW, Jones T, Day S, Myers L, Eun-Hyung Lee F, Crowe JE. 2020. Rapid isolation and profiling of a diverse panel of human monoclonal antibodies targeting the SARS-CoV-2 spike protein. Nat Med 26:1422–1427.

21. FDA. 2021. Coronavirus (COVID-19) Update: FDA Revokes Emergency Use Authorization for Monoclonal Antibody Bamlanivimab | FDA. FDA NEWS RELEASE.

22. CDC. 2021. SARS-CoV-2 Variant Classifications and Definitions.

23. McCallum M, De Marco A, Lempp FA, Tortorici MA, Pinto D, Walls AC, Beltramello M, Chen A, Liu Z, Zatta F, Zepeda S, di Iulio J, Bowen JE, Montiel-Ruiz M, Zhou J, Rosen LE, Bianchi S, Guarino B, Fregni CS, Abdelnabi R, Veesler D. 2021. N-terminal domain antigenic mapping reveals a site of vulnerability for SARS-CoV-2. Cell 184:2332–2347.e16.

24. Suryadevara N, Shrihari S, Gilchuk P, VanBlargan LA, Binshtein E, Zost SJ, Nargi RS, Sutton RE, Winkler ES, Chen EC, Fouch ME, Davidson E, Doranz BJ, Chen RE, Shi P-Y, Carnahan RH, Thackray LB, Diamond MS, Crowe JE. 2021. Neutralizing and protective human monoclonal antibodies recognizing the N-terminal domain of the SARS-CoV-2 spike protein. Cell 184:2316–2331.e15.

25. Chi X, Yan R, Zhang J, Zhang G, Zhang Y, Hao M, Zhang Z, Fan P, Dong Y, Yang Y, Chen Z, Guo Y, Zhang J, Li Y, Song X, Chen Y, Xia L, Fu L, Hou L, Xu J, Chen W. 2020. A neutralizing human antibody binds to the N-terminal domain of the Spike protein of SARS-CoV-2. Science 369:650–655.

26. Xu C, Wang Y, Liu C, Zhang C, Han W, Hong X, Wang Y, Hong Q, Wang S, Zhao Q, Wang Y, Yang Y, Chen K, Zheng W, Kong L, Wang F, Zuo Q, Huang Z, Cong Y. 2021. Conformational dynamics of SARS-CoV-2 trimeric spike glycoprotein in complex with receptor ACE2 revealed by cryo-EM. Sci Adv 7.

27. Cerutti G, Guo Y, Zhou T, Gorman J, Lee M, Rapp M, Reddem ER, Yu J, Bahna F, Bimela J, Huang Y, Katsamba PS, Liu L, Nair MS, Rawi R, Olia AS, Wang P, Zhang B, Chuang G-Y, Ho DD, Shapiro L. 2021. Potent SARS-CoV-2 neutralizing antibodies directed against spike N-terminal domain target a single supersite. Cell Host Microbe 29:819–833.e7.

28. Wang N, Sun Y, Feng R, Wang Y, Guo Y, Zhang L, Deng Y-Q, Wang L, Cui Z, Cao L, Zhang Y-J, Li W, Zhu F-C, Qin C-F, Wang X. 2020. Structure-based development of human antibody cocktails against SARS-CoV-2. Cell Res 31:101–103.

29. Wec AZ, Bornholdt ZA, He S, Herbert AS, Goodwin E, Wirchnianski AS, Gunn BM, Zhang Z, Zhu W, Liu G, Abelson DM, Moyer CL, Jangra RK, James RM, Bakken RR, Bohorova N, Bohorov O, Kim DH, Pauly MH, Velasco J, Chandran K. 2019. Development of a Human Antibody Cocktail that Deploys Multiple Functions to Confer Pan-Ebolavirus Protection. Cell Host Microbe 25:39–48.e5.

30. Bornholdt ZA, Herbert AS, Mire CE, He S, Cross RW, Wec AZ, Abelson DM, Geisbert JB, James RM, Rahim MN, Zhu W, Borisevich V, Banadyga L, Gunn BM, Agans KN, Wirchnianski AS, Goodwin E, Tierney K, Shestowsky WS, Bohorov O, Dye JM. 2019. A Two-Antibody Pan-Ebolavirus Cocktail Confers Broad Therapeutic Protection in Ferrets and Nonhuman Primates. Cell Host Microbe 25:49–58.e5.

31. Dieterle ME, Haslwanter D, Bortz RH, Wirchnianski AS, Lasso G, Vergnolle O, Abbasi SA, Fels JM, Laudermilch E, Florez C, Mengotto A, Kimmel D, Malonis RJ, Georgiev G, Quiroz J, Barnhill J, Pirofski L-A, Daily JP, Dye JM, Lai JR, Jangra RK. 2020. A Replication-Competent Vesicular Stomatitis Virus for Studies of SARS-CoV-2 Spike-Mediated Cell Entry and Its Inhibition. Cell Host Microbe 28:486–496.e6.

32. Weisblum Y, Schmidt F, Zhang F, DaSilva J, Poston D, Lorenzi JC, Muecksch F, Rutkowska M, Hoffmann H-H, Michailidis E, Gaebler C, Agudelo M, Cho A, Wang Z, Gazumyan A, Cipolla M, Luchsinger L, Hillyer CD, Caskey M, Robbiani DF, Bieniasz PD. 2020. Escape from neutralizing antibodies by SARS-CoV-2 spike protein variants. elife 9.

33. Andreano E, Piccini G, Licastro D, Casalino L, Johnson NV, Paciello I, Monego SD, Pantano E, Manganaro N, Manenti A, Manna R, Casa E, Hyseni I, Benincasa L, Montomoli E, Amaro RE, McLellan JS, Rappuoli R. 2020. SARS-CoV-2 escape in vitro from a highly neutralizing COVID-19 convalescent plasma. BioRxiv https://doi.org/10.1101/2020.12.28.424451.

34. Sakharkar M, Rappazzo CG, Wieland-Alter WF, Hsieh C-L, Wrapp D, Esterman ES, Kaku CI, Wec AZ, Geoghegan JC, McLellan JS, Connor RI, Wright PF, Walker LM. 2021. Prolonged evolution of the human B cell response to SARS-CoV-2 infection. Sci Immunol 6.

35. Gaebler C, Wang Z, Lorenzi JCC, Muecksch F, Finkin S, Tokuyama M, Cho A, Jankovic M, Schaefer-Babajew D, Oliveira TY, Cipolla M, Viant C, Barnes CO, Bram Y, Breton G, Hägglöf T, Mendoza P, Hurley A, Turroja M, Gordon K, Nussenzweig MC. 2021. Evolution of antibody immunity to SARS-CoV-2. Nature 591:639–644.

36. Hartley GE, Edwards ESJ, Aui PM, Varese N, Stojanovic S, McMahon J, Peleg AY, Boo I, Drummer HE, Hogarth PM, O’Hehir RE, van Zelm MC. 2020. Rapid generation of durable B cell memory to SARS-CoV-2 spike and nucleocapsid proteins in COVID-19 and convalescence. Sci Immunol 5.

37. Briney B, Inderbitzin A, Joyce C, Burton DR. 2019. Commonality despite exceptional diversity in the baseline human antibody repertoire. Nature 566:393–397.

38. Jones BE, Brown-Augsburger PL, Corbett KS, Westendorf K, Davies J, Cujec TP, Wiethoff CM, Blackbourne JL, Heinz BA, Foster D, Higgs RE, Balasubramaniam D, Wang L, Zhang Y, Yang ES, Bidshahri R, Kraft L, Hwang Y, Žentelis S, Jepson KR, Falconer E. 2021. The neutralizing antibody, LY-CoV555, protects against SARS-CoV-2 infection in nonhuman primates. Sci Transl Med 13.

39. FDA. 2021. EMERGENCY USE AUTHORIZATION (EUA) OF BAMLANIVIMAB AND ETESEVIMAB AUTHORIZED USE.

40. FDA. 2021. FACT SHEET FOR HEALTH CARE PROVIDERS EMERGENCY USE AUTHORIZATION (EUA) OF REGEN-COVTM (casirivimab with imdevimab).

41. Voss WN, Hou YJ, Johnson NV, Delidakis G, Kim JE, Javanmardi K, Horton AP, Bartzoka F, Paresi CJ, Tanno Y, Chou C-W, Abbasi SA, Pickens W, George K, Boutz DR, Towers DM, McDaniel JR, Billick D, Goike J, Rowe L, Ippolito GC. 2021. Prevalent, protective, and convergent IgG recognition of SARS-CoV-2 non-RBD spike epitopes. Science https://doi.org/10.1126/science.abg5268.

42. Chou T-C, Talalay P. 1984. Quantitative analysis of dose-effect relationships: the combined effects of multiple drugs or enzyme inhibitors. Adv Enzyme Regul 22:27–55.

43. Zhou H, Chen Y, Zhang S, Niu P, Qin K, Jia W, Huang B, Zhang S, Lan J, Zhang L, Tan W, Wang X. 2019. Structural definition of a neutralization epitope on the N-terminal domain of MERS-CoV spike glycoprotein. Nat Commun 10:3068.

44. Deng X, Garcia-Knight MA, Khalid MM, Servellita V, Wang C, Morris MK, Sotomayor-González A, Glasner DR, Reyes KR, Gliwa AS, Reddy NP, Sanchez San Martin C, Federman S, Cheng J, Balcerek J, Taylor J, Streithorst JA, Miller S, Kumar GR, Sreekumar B, Chiu CY. 2021. Transmission, infectivity, and antibody neutralization of an emerging SARS-CoV-2 variant in California carrying a L452R spike protein mutation. medRxiv https://doi.org/10.1101/2021.03.07.21252647.

45. CDC. 2021. CDC 2019-Novel Coronavirus (2019-nCoV) Real-Time RT-PCR Diagnostic Panel.

46. Daniel Gietz R, Woods RA. 2002. Transformation of yeast by lithium acetate/single-stranded carrier DNA/polyethylene glycol method, p. 87–96. In Guide to Yeast Genetics and Molecular and Cell Biology - Part B. Elsevier.

47. Rappazzo CG, Tse LV, Kaku CI, Wrapp D, Sakharkar M, Huang D, Deveau LM, Yockachonis TJ, Herbert AS, Battles MB, O’Brien CM, Brown ME, Geoghegan JC, Belk J, Peng L, Yang L, Hou Y, Scobey TD, Burton DR, Nemazee D, Walker LM. 2021. Broad and potent activity against SARS-like viruses by an engineered human monoclonal antibody. Science 371:823–829.

